# Mouse model and human patient data suggest critical roles for Pten and p53 in suppressing POLE mutant tumor development

**DOI:** 10.1101/2021.10.10.463847

**Authors:** Vivian S. Park, Meijuan J.S. Sun, Wesley D. Frey, Leonard G. Williams, Karl P. Hodel, Juliet D. Strauss, Sydney J. Wellens, James G. Jackson, Zachary F. Pursell

**Author notes:** To whom correspondence should be addressed: Zachary F. Pursell.

## Abstract

Mutations in the exonuclease domain of *POLE* are associated with tumors harboring very high mutation burdens. The mechanisms linking this significant mutation accumulation and tumor development remain poorly understood. *Pole*^+/P286R^;*Trp53*^+/−^ mice showed accelerated cancer mortality compared to *Pole*^+/P286R^;*Trp53*^+/+^ mice. Cells from *Pole*^+/P286R^ mice showed increased p53 activation, and subsequent loss of p53 permitted rapid growth, implicating canonical p53 loss of heterozygosity in POLE mutant tumor growth. Somewhat surprisingly, however, p53 status had no effect on tumor mutation burden or single base substitution signatures in *POLE* mutant tumors from mice or humans. Pten has important roles in maintaining genome stability. We find that *PTEN* mutations are highly enriched in human POLE mutant tumors, including many in POLE signature contexts. One such signature mutation, *PTEN-F341V*, was previously shown in a mouse model to specifically decrease nuclear Pten and lead to increased DNA damage. We found tumors in *Pole*^+/P286R^ mice that spontaneously acquired *Pten*^*F341V*^ mutations and were associated with significantly reduced nuclear Pten and elevated DNA damage. Taken together with recent published work, our results support the idea that POLE-mediated hypermutagenesis is necessary, but not entirely sufficient, for tumorigenesis. Disabling surveillance of nuclear DNA damage is a likely sufficient factor.

## INTRODUCTION

Tumors with Pol ɛ exonuclease domain mutations are associated with incredibly high mutation burdens exceeding >10 mutations per Megabase (mut/Mb) and unique mutation signatures that distinguish them from other cancers. These signatures include T<u>C</u>T>T<u>A</u>T transversions, T<u>C</u>G>T<u>T</u>G transitions and T<u>T</u>T>T<u>G</u>T transversions, designated as single base substitution (SBS) signatures 10a, 10b and 28, respectively ^1^. These signatures are significantly elevated in POLE mutant tumors from both humans ^1,2^ and mice ^3^. An additional signature, SBS14, is found in human POLE tumors that also show evidence of microsatellite instability and inactivation of one or more mismatch repair factors. SBS14 is dominated by N<u>C</u>T>N<u>A</u>T transversions, with N representing any base (SBS 14) ^1,4^. However, the mechanism(s) by which mutations in *POLE* lead to tumorigenesis are not well understood. We used our established germline *Pole*-P286R mouse, which models the most recurrent human *POLE* mutation ^2,5–7^, to 1) investigate how *POLE* mutations drive tumorigenesis *in vivo* and 2) determine the extent to which tumor development is driven by mutagenesis *per se* or by non-mutagenic processes.

We previously reported the curious finding that *Pole*^*S459F/S459F*^ mice were viable and developed to adulthood, yet *Pole*^*P286R/P286R*^ mice were not viable, even in the same background strain ^3^. This was somewhat surprising because the exonuclease activity of the human *POLE*-S459F enzyme is 100-fold lower than *POLE*-P286R in *in vitro* studies ^3^. Since mutation accumulation and mutation spectra have been shown to be influenced by the nature of the mutant *POLE* allele ^8–10^, it follows that these and other factors also likely contribute to normal embryonic development.

To further understand what factors might contribute to the high mutagenesis observed in murine *POLE* mutant tumors and to the *Pole*^*P286R/P286R*^ embryonic lethality phenotype, we bred *Pole*^+/*P286R*^ mice with *Trp53*^+/−^ mice ^11^. The rationale guiding this cross was driven by several lines of evidence suggesting that p53 activation is a downstream target of POLE dysfunction. Bellelli et al. found that *Pole4*^−/−^ mice, which lack a non-catalytic Pol ε holoenzyme subunit, had developmental abnormalities and increased tumorigenesis ^12^. *Pole4*^−/−^ was embryonic lethal in a C57BL/6J background, but this lethality was rescued when p53 was inactivated. They also reported that homozygous *Pole4*^−/−^ embryos had significantly increased p53 protein levels compared to wild-type and heterozygous embryos ^12^. We have previously reported that mRNA levels of p21 and PUMA are elevated in human cell lines engineered to express mutant *POLE* alleles ^9^. Additionally, human *POLE*-mutated tumors are associated with enrichment in mutations in major tumor suppressor genes, including *TP53* ^13,14^. Therefore, we reasoned that *Pole*^P286R/P286R^ and perhaps *Pole*^+/P286R^ survival might be enhanced in the absence of functional p53.

While the sheer number of mutated genes in *POLE* mutated tumors has made identifying tumor driving mutations difficult, a number of studies have made attempts. Enrichment of *TP53* mutations in POLE mutant tumors, in particular the R231X nonsense mutation, was noted in several studies ^14,15^. A more careful subsequent look at p53 mutations in POLE mutant tumors found that p53-R213X was significantly elevated specifically in CRC tumors with the *POLE*^*P286R*,^ mutant allele ^13^. Neither endometrial tumors nor those with the *POLE*^*V411L*^ mutant allele showed enrichment in the p53-R213X mutation. Almost half of mouse endometrial tumors formed due to conditional knockin of *Pole*^+/P286R^ in uterine tissue stained positively for p53, though the nature of the mutant alleles was not examined further ^16^. Another candidate that has been reported in addition to p53 in human POLE mutant tumors is the tumor suppressor PTEN ^17,18^, though its association with POLE mutations was scored as indeterminate.

In the current study we looked at how mutations in *Trp53* affected growth and mutagenesis in cells and tumors from *Pole*^P286R^ mice. We then used information from human tumor datasets to investigate the relationship between mutagenesis and tumorigenesis in POLE mutant tumors. We found that loss of one *Trp53* allele accelerated tumorigenesis without affecting mutation burden or mutation signature. Similar to previous observations in engineered human cell lines, p53 was activated in mouse cells with mutant alleles of *Pole*. While *TP53* driver mutations were not significantly enriched in human *POLE* mutant tumors, we found enriched and recurrent *PTEN* mutations occurring in mutant POLE signature contexts. Analyzing a human endometrial tumor proteogenomic dataset revealed that POLE-dependent mutations correlated with reduced Pten protein levels while p53 levels were unchanged. Further, we found spontaneously occurring *Pten*^F341V^ mutations in two separate *Pole*^P286R^ mouse tumors that displayed loss of nuclear Pten staining. Previous studies using a germline *Pten*^F341V^ mouse model have shown this allele drives increased tumorigenesis coupled with decreased genome stability. Taken together we propose that loss of p53 or loss of nuclear Pten can facilitate POLE mutant tumor development, likely through loss of DNA damage surveillance.

## RESULTS

### Loss of p53 does not suppress Pole^P286R/P286R^ embryonic lethality

Loss of p53 function has previously been implicated in POLE mutant tumor development in humans ^13,14^ and in mice ^16^. We have also shown that p53 is activated in human cancer cell lines engineered to express cancer mutant alleles of *POLE* ^9^. We previously characterized the effects of the *Pole*-P286R mutation on tumor development and mutagenesis in a mouse model ^3^. We noted that *Pole*^P286R/P286R^ was likely embryonic lethal due to lack of any observed viable offspring. Inactivation of p53 has been used to rescue embryonic lethality for many mouse mutant alleles, including *Pole4*^−/−^ ^12,19^.

We set out to test the effects of p53 loss on POLE mutant tumor development by inactivating *Trp53* in our *Pole*-P286R mice. We crossed *Pole*^+/P286R^ and *Trp53*^−/−^ mice with one another to generate double heterozygous, *Pole*^+/P286R^*;Trp53*^+/−^ mice. Breeding double heterozygous mice (*Pole*^+/P286R^*;Trp53*^+/−^ × *Pole*^+/P286R^*;Trp53*^+/−^) produced one *Pole*^P286R/P286R^;*Trp53*^+/−^ pup out of 38 live births; however, this offspring failed to thrive and died one day post-utero (Supplementary Table 1). We also observed one *Pole*^P286R/P286R^ embryo *in utero* out of one litter genotyped at approximately embryonic day 13.5-14.5 (*Pole*^P286R/P286R^*;Trp53*^+/−^, red arrow, Supplementary Figure 1A). Compared to littermates, the *Pole*^*P286R/P286R*^ embryo was noticeably underdeveloped and likely would not have survived to live birth. In addition, we observed one *Pole*^+/*P286R*^*;Trp53*^−/−^ embryo, which was also comparatively smaller than its littermates. The lack of expected Mendelian ratio of alleles in viable offspring indicates that loss of p53 does not suppress lethality in developing *Pole*^*P286R/P286R*^ embryos.

While breeding *Pole*^+/P286R^*;Trp53*^+/−^ mice, we also observed that they displayed accelerated mortality compared to *Pole*^+/P286R^*;Trp53*^+/+^ mice. In order to directly test the effect of *Trp53* status on survival of *Pole*^+/P286R^ we set up a survival cohort of double heterozygous breeding pairs (*Pole*^+/*P286R*^;*Trp53*^+/−^ × *Pole*^+/*P286R*^;*Trp53*^+/−^) to generate sufficient numbers of littermates of all possible genotypes. Survival of *Pole*^+/P286R^;*Trp53*^+/−^ mice was significantly decreased compared to *Pole*^+/+^*;Trp53*^+/+^ mice, with survival ranging from 2.0 to 9.1 months and median survival at 4 months (Figure 1A, p=0.03). No significant difference in survival of male vs. female mice within the same genotype was observed. Tumors from *Pole*^+/P286R^*;Trp53*^+/−^ mice showed similar lymphoid tissue specificity to those from *Pole*^+/P286R^*;Trp53*^+/+^ mice (Supp. Table 2).

**Figure 1.**
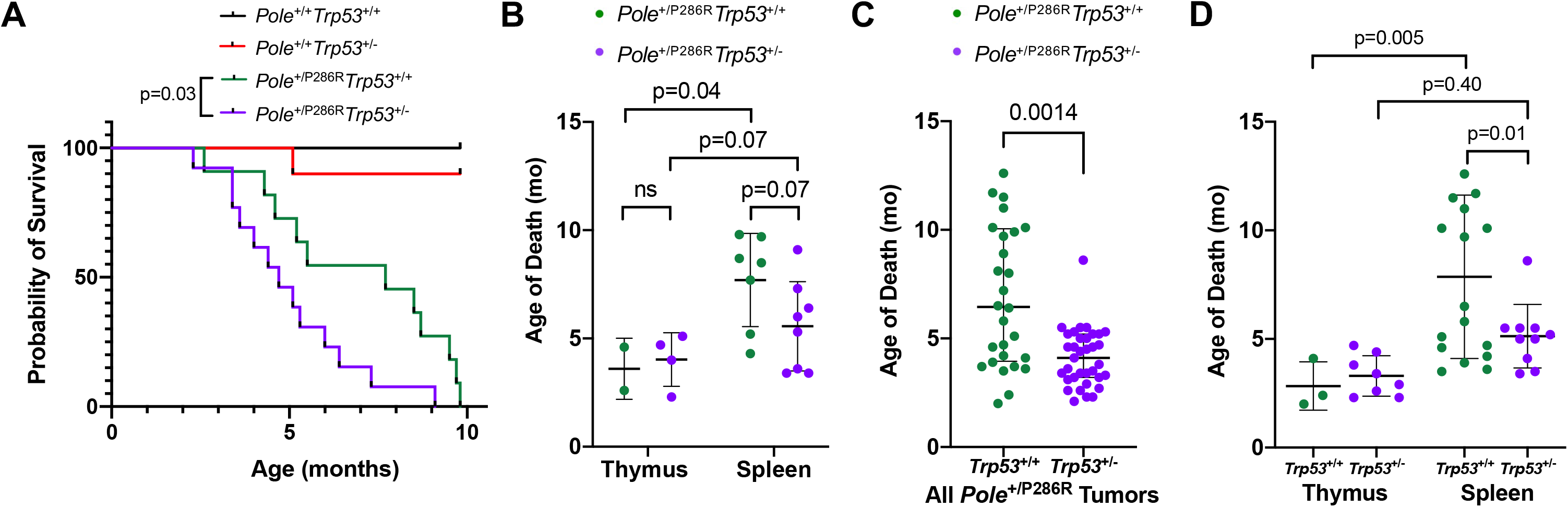
Loss of *Trp53* allele accelerates *Pole*-dependent cancer mortality. **(A)** Kaplan-Meier survival estimate of *Pole*^+/*P286R*^;*Trp53*^+/−^ mice compared to *Pole*^+/*P286R*^;*Trp53*^+/+^ mice. All mice were products of *Pole*^+/*P286R*^;*Trp53*^+/−^ × *Pole*^+/*P286R*^;*Trp53*^+/−^ cross. *Pole*^+/+^;Trp53^+/+^ (n = 8); *Pole*^+/+^;Trp53^+/−^ (n = 10); *Pole*^+/*P286R*^;*Trp53*^+/+^ (n = 11); *Pole*^+/*P286R*^;*Trp53*^+/−^ (n = 13). **(B)** Cohort mean age of death stratified by thymic and disseminated (spleen) lymphomas. Mean age: 3.6, 4.0, 7.7 and 5.6 months, respectively. For comparisons of the means between two groups, unpaired two-tailed t-test and Welch’s correction for nonequal SD was used. **(C)** Mean age of death for *Pole*^+/*P286R*^ and *Pole*^+/*P286R*^;*Trp53*^+/−^ mice from all tumors (expanded to mice beyond cohort in (A and B)), 7.0 to 4.1 months, respectively. **(D)** Mean age of death for *Pole*^+/*P286R*^ and mice (expanded to mice beyond cohort (shown in A and B) stratified by thymic and disseminated (spleen) lymphomas. Mean age: 2.8, 3.3, 7.9 and 5.1 months, respectively. For comparisons of the means between two groups, unpaired two-tailed t-test and Welch’s correction for nonequal SD was used.

While overall cancer mortality was accelerated by loss of one allele of p53, the effect was not the same across tissue types. *Pole*^+/P286R^ mice with thymic lymphomas died rapidly regardless of germline *Trp53* status (Fig. 1B). Occurrence of disseminated lymphomas in the spleen, lymph nodes and liver was slightly accelerated in *Pole*^+/P286R^*;Trp53*^+/−^ mice compared to *Pole*^+/P286R^*;Trp53*^+/+^ mice (Fig. 1B, 5.6 months vs 7.7 months, respectively), though this difference was not significant in the survival cohort. When we looked more broadly at each of the first 63 mice (*Pole*^+/*P286R*^;*Trp53*^+/+^ (n = 28) and *Pole*^+/*P286R*^;*Trp53*^+/−^(n = 35)) generated from the beginning of this colony and compared to *Pole*^+/*P286R*^ mice from our previous study ^3^, overall tumor mortality was accelerated in *Pole*^+/P286R^*;Trp53*^+/−^ mice (Fig. 1C). Interestingly, this difference was entirely due to more rapid development of splenic lymphomas, as thymic lymphoma development remained independent of *Trp53* genotype (Fig. 1D). Based on this expanded data, loss of one allele of *Trp53* significantly decreased survival, likely by accelerating disseminated tumor mortality.

### p53 activation in cells with mutant Pole restrains immortalization

We used mouse embryonic fibroblasts (MEFs) to address the effects of *Pole*-P286R on p53 in untransformed mouse cells. We found that time to immortalization was dependent on both *Pole* and *Trp53* status (Fig. 2A). *Pole*^+/*P286R*^;*Trp53*^−/−^ MEFs did not plateau and continued steady population doubling levels throughout the experiment. Population doubling levels (PDLs) for *Pole*^+/*P286R*^;*Trp53*^+/−^ MEFs slowed through 21 days, followed by rapid proliferation, indicating likely immortalization through inactivation of the remaining *Trp53* allele. Consistent with NIH 3T3 mouse embryo cells in culture ^20^, PDLs for both *Pole*^+/P286R^*;Trp53*^+/+^ and *Pole*^+/+^*;Trp53*^+/+^ MEFs plateaued after about 9 days and remained so after more than 30 days of passaging.

**Figure 2.**
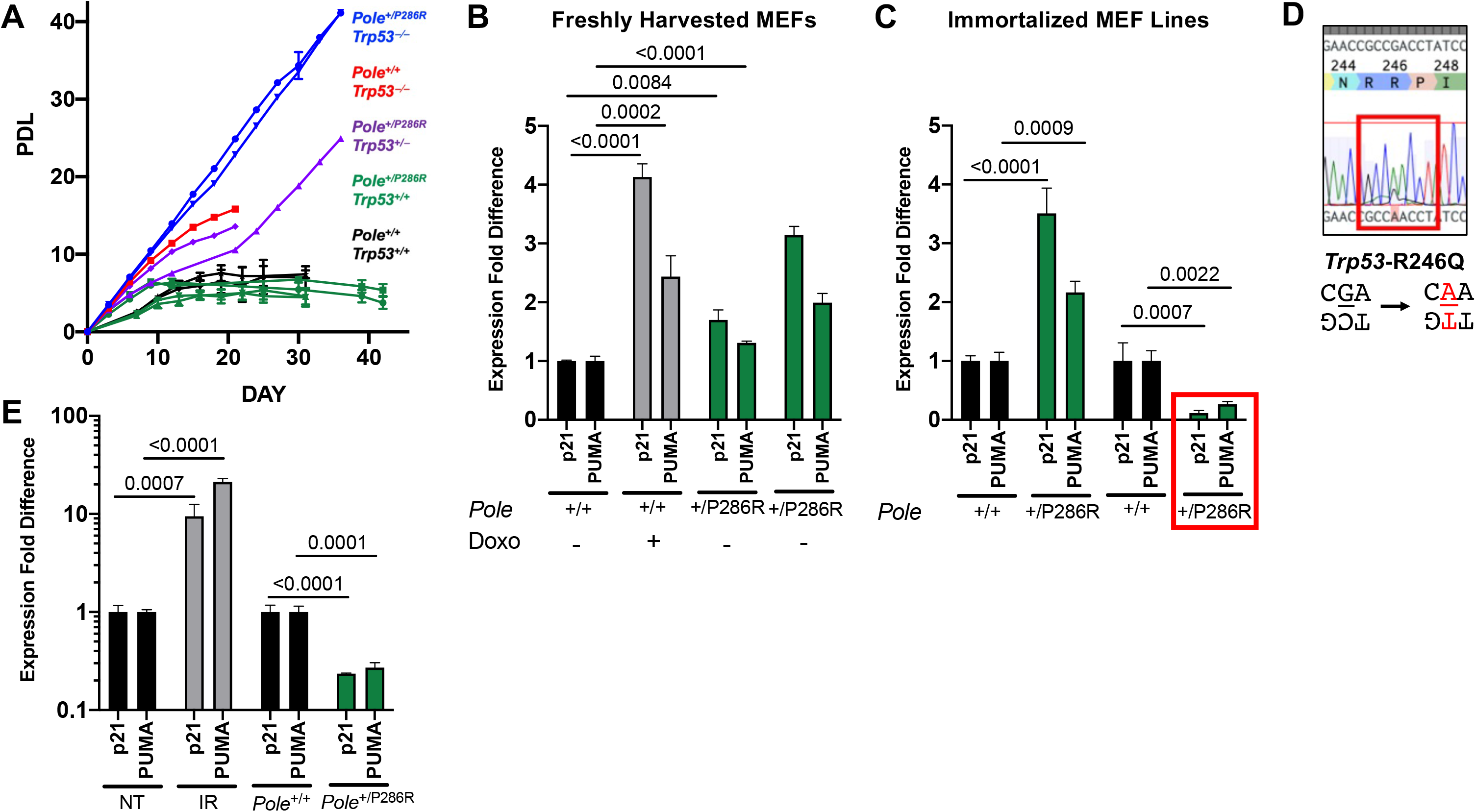
Activation of p53 target expression in *Pole*^+/*P286R*^ MEFs and immortalized mouse cell lines. **(A)** Population doubling levels (PDL) of MEFs passaged according to NIH 3T3 protocol double heterozygous crosses. Genotypes include: *Pole*^+/+^;*Trp53*^+/+^, *Pole*^+/+^;*Trp53*^−/−^, *Pole*^+/*P286R*^;*Trp53*^+/+^, *Pole*^+/*P286R*^;*Trp53*^+/−^, *Pole*^+/*P286R*^;*Trp53*^−/−^. **(B)** qPCR expression of p21 and PUMA in freshly derived *Pole*^+/*P286R*^ MEF cell lines. 4226 cell line, created from p53 wild-type MMVT-*Wnt1* tumor cells, non-treated (NT) and doxorubicin-treated (Doxo) cells included as positive control for p53 activation. **(C)** qPCR expression of p21 and PUMA in immortalized *Pole*^+/*P286R*^ and *POLE*^+/*P286R*^ cells lines. **(D)** Cell line with significantly decreased p53 activation had mutation in DNA binding domain of *Trp53* verified via Sanger sequencing. **(E)** qPCR expression of p21 and PUMA in *Pole*^+/*P286R*^ thymic tumor compared to normal tissue determined by qPCR. Non-treated (NT) and irradiated (IR) thymus included as positive control for p53 activation.

Consistent with the proliferation assays, *Cdkn1a* and *Bbc3* expression was elevated in MEF cells freshly harvested and plated from *Pole*^+/P286R^*;Trp53*^+/+^ embryos (Fig. 2B). As in our human cell line models ^9^, activation of p53 targets was also significantly elevated in immortalized *Pole*^+/*P286R*^ mouse cell lines when compared to *Pole*^+/+^ cell lines (Fig. 2C, left). However, we noted that one spontaneously immortalized *Pole*^+/*P286R*^ cell line had significantly reduced p53 activation (Fig. 2C, right). Sequencing the DNA encoding the DNA binding domain of p53 from these cells showed that they acquired a p53-R246Q mutation (Fig. 2D), which has a known dominant negative function in mice ^21,22^. Interestingly, the R246Q mutation is due to a C>T-T<u>C</u>G POLE signature 10b mutation on the noncoding strand (Fig. 2D), indicating that mutant *Pole* was likely directly responsible for p53 inactivation. Thymic tumors also had significantly reduced levels of p53 activation compared to normal thymus tissue, presumably allowing uncontrolled growth (Fig. 2E). Taken together, these data establish, in two model systems, a direct link between mutant POLE-mediated mutagenesis and p53 activation and selective pressure to mutate the *Trp53* to enable immortalization and transformation.

### Loss of p53 loss has minimal impact on POLE-mediated tumor mutation accumulation

Since we had previously showed that *Pole*^+/P286R^ could accelerate cancer mortality in mice and was associated with hypermutant POLE mutagenesis phenotype, we investigated the effect of p53 loss on mutagenesis. We selected four tumors from *Pole*^+/*P286R*^;*Trp53*^+/−^ mice and performed WES. Each tumor had mutations in the remaining Trp53 allele, including one known loss of function mutation (R210Q ^23–25^), one loss of function frameshift mutation (Q97fs) [cite Giac], one mutation that was shown to occur multiple times in human breast cancer patients (L194R ^26^) and one charge reversal that was predicted to reduce function in a saturation mutagenesis screen (E14K ^23^) (Fig. 3A). These results indicate that tumors in *Pole*^+/*P286R*^;*Trp53*^+/−^ mice demonstrate selective pressure for inactivating p53 in the development of tumors driven by *Pole* mutation. All four mutations occurred in POLE signature sequence contexts, strongly suggesting that the mutator polymerase first made the DNA synthesis error, and cells with this p53 mutation subsequently proliferated and expanded during tumor development.

**Figure 3.**
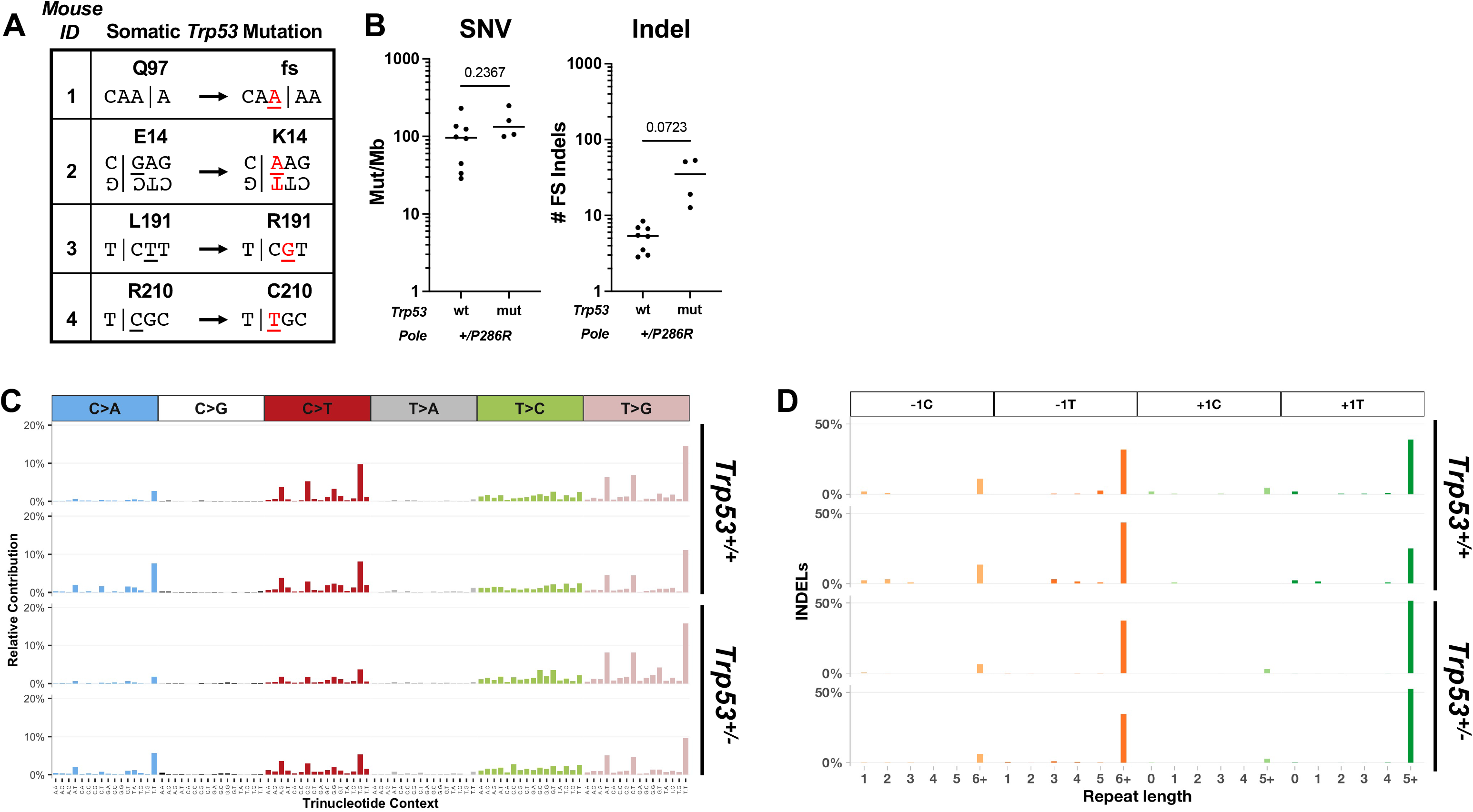
*Pole*^+/*P286R*^ tumors display high mutation burden and POLE signatures idenpendent of p53 status. **(A)** Spontaneous somatic *Trp53* mutations arise in all tumors from germline *Pole*^+/*P286R*^;*Trp53*^+/−^ mice. **(B)** Single nucleotide variants (SNV) and insertion and deletion frameshift mutations (Indels) in tumors from *Pole*^+/*P286R*^;*Trp53*^+/+^ and *Pole*^+/*P286R*^;*Trp53*^+/−^ mice. For comparisons of the means between two groups, unpaired two-tailed t-test and Welch’s correction for nonequal SD were used. Mutation spectra of SNVs **(C)** and indels **(D)** from two *Pole*^+/*P286R*^;*Trp53*^+/+^ thymic tumors (top) and two *Pole*^+/*P286R*^;*Trp53*^+/−^ splenic tumors (bottom) are shown.

Somewhat surprisingly, however, p53 status did not affect overall base pair substitution in tumors from *Pole*^+/P286*R*^ mice (Fig. 3B, left). We did notice that insertion/deletion (indel) mutations were elevated in tumors from *Pole*^+/P286R^;*Trp53*^+/−^ (Fig. 3B, right), but this difference was not significant. All POLE mutation signatures (SBS 10a, 28 and 10b) were also present in tumors from *Pole*^+/*P286R*^;*Trp53*^+/−^ mice (Fig. 3C) ^3^, as were the more recently described +A insertions into runs of A nucleotides, particularly those ≥ 5 nt ^27^ (Supp. Fig. 2).

Using the TCGA endometrial and colorectal tumor datasets, we examined how tumor mutation burden and POLE signature mutations in human tumors are affected by the presence of TP53 mutations. We stratified all tumors with *POLE* driver mutations ^9,28^ into those with known *TP53* driver mutations (*TP53* mut, as defined by cBIOPORTAL ^29^) and those without known driver mutations (*TP53* wt). The *TP53* wt set may contain mutations of unknown significance. Consistent with our data in mice, incidence of SNVs and indels did not differ between *POLE* mutant tumors that were *TP53* mutant or *TP53* wild type (Fig. 4A).

**Figure 4.**
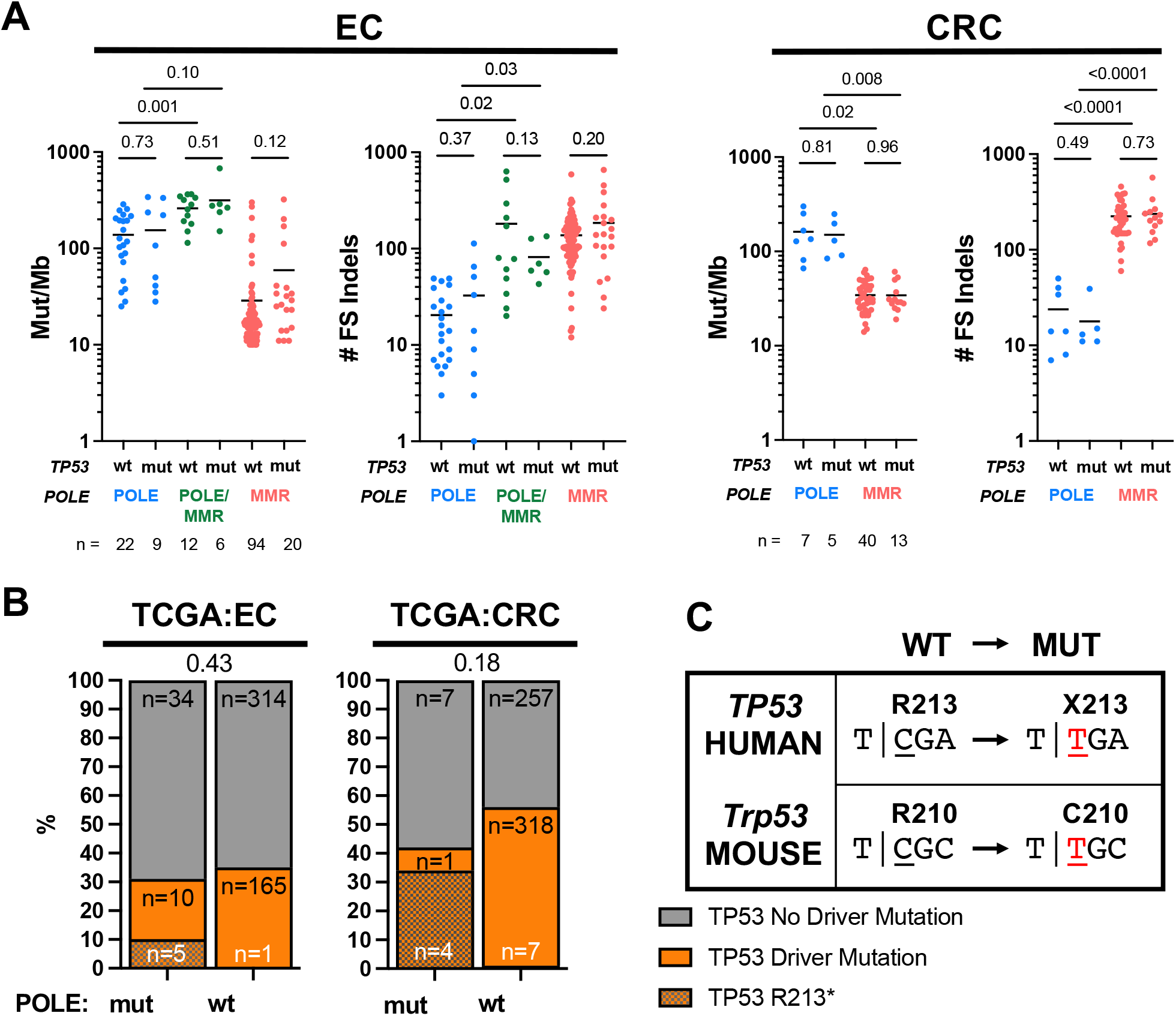
Driver mutations in *TP53* do not affect mutagenesis in TCGA endometrial (EC) and colorectal (CRC) cancers. **(A)** Mutation burden of SNVs (Mut/Mb) and number of insertions/deletions (Indels), categorized by POLE, POLE/MMR and MMR SBS signature status. Comparison is stratified by *TP53* driver mutation status. p-values indicated and determined by ANOVA. **(B)** Frequencies of *TP53* driver mutations in endometrial and colorectal tumors with mutations in POLE are no different than in POLE wild type tumors. Fisher’s exact test was performed comparing all *TP53* driver mutations, p-values indicated above stacked bars. **(C)** The single POLE mutation hotspot in *TP53* from human tumors is not a POLE hotspot in mouse *Trp53*.

Mutagenesis in POLE mutant tumors and cell lines has been shown to be a composite of several different signatures that depends in part on the mutant allele of *POLE* and whether mismatch repair (MMR) is functional or not ^4,9^. We asked whether p53 status differentially affected mutagenesis based on POLE and MMR status by categorizing tumors based on mutational signatures: POLE (SBS 10a, 10b or 28 in **blue**), POLE/MMR (SBS 14 in **green**) and MMR ^30^ (SBS 6, 15, 20, 21, 26 or 44 in **red**). *TP53* driver mutation status did not affect SNVs or indels (Fig. 4A) in POLE, POLE/MMR and MMR tumors. Taken together, these results suggest that the acceleration of tumor development in POLE mutant tumors that have lost p53 function is not due to simple increased or altered pattern of mutagenesis.

### Lack of p53 mutation enrichment in human POLE mutant tumors

Previous analyses of cancer mutations showed enrichment for the p53 R213X mutation in POLE mutant tumors ^13,14^. In order to examine all *TP53* driver mutations in POLE mutant tumors, we first defined *POLE*-mutated tumors (POLE mut) by two criteria as described: TMB >10 mut/Mb and the presence of SBS 10a, 10b and 28 ^9^. Tumors not satisfying both criteria were considered wild type for POLE (POLE wt). We limited this analysis to TCGA uterine and colorectal tumors (UCEC and CRC, respectively) since they account for 90% (61 of 68) of all POLE mutant tumors in the TCGA database. Samples in cBioportal were further classified as having a driver mutation in *TP53* or not ^29,31^ based on OncoKB and CancerHotspots databases of curated cancer mutations known to have functional and clinical relevance in model organisms and patients as described ^32,33^. One UCEC patient was not included as no specific gene mutation data were available in cBioPortal (TCGA-EY-A1G8).

Surprisingly, the overall frequency of *TP53* driver mutations was no different in *POLE*-mutated and *POLE* wild-type tumors in the TCGA UCEC and CRC cohorts (Fig. 4B). R213X mutations were clearly enriched in POLE mutant tumors, accounting for 33% and 80% of the *TP53* driver set of mutations in UCEC and CRC tumors, respectively (Fig. 4B). As others have noted, the R213 codon in human *TP53* combined with the 3′ base of the immediate upstream codon results in the POLE signature hotspot, T <u>C</u>GA. POLE-dependent mutagenesis then generates an opal TGA nonsense mutation encoding R213X (Fig. 4C). We note that the absence of p53 R213X mutations in our mouse tumor samples is easily explained by alternate arginine codon usage (CGC) in the mouse genome eliminating the POLE signature context.

### Enrichment of mutations in other tumor suppressors in The Cancer Genome Atlas human tumors and mice

Since p53 driver mutations were not enriched in *POLE*-mutated tumors (Fig. 4B), we examined the relationship between mutations in *POLE* and in other commonly mutated cancer genes and pathways. Driver mutations in *PTEN*, a negative regulator of the PI3K/AKT pathway, were significantly enriched in both UCEC and CRC (Fig. 5A). This enrichment had been previously noted in two endometrial tumor studies ^17,34^, but not in TCGA. *PTEN* driver mutations are well known to be significantly enriched in UCEC tumors ^35^ and indeed we observed 56% of all TCGA uterine tumors have such mutations. We found an even further increase in POLE mutant tumors, with almost all (94%) harboring a *PTEN* driver mutation (Fig. 5A, p<0.0001, Fisher’s exact test). While the absolute number of POLE mutant tumors is lower, this enrichment still occurred in CRC tumors (Fig. 5A, p<0.0001, Fisher’s exact test). We validated this *PTEN* driver mutation enrichment in POLE mutant tumors in two separate endometrial tumor cohorts: the CPTAC proteogenomic dataset (Fig. 5B) and the MSK-IMPACTS endometrial dataset (Supp. Fig. 3). Mutations in *POLE* also occur in tumors other than uterine or colorectal, albeit quite infrequently. However, even in these disparate tumors we still found *PTEN* mutation enrichment in POLE mutant tumors (Fig. 5B), further supporting the idea that these pathways interact.

**Figure 5.**
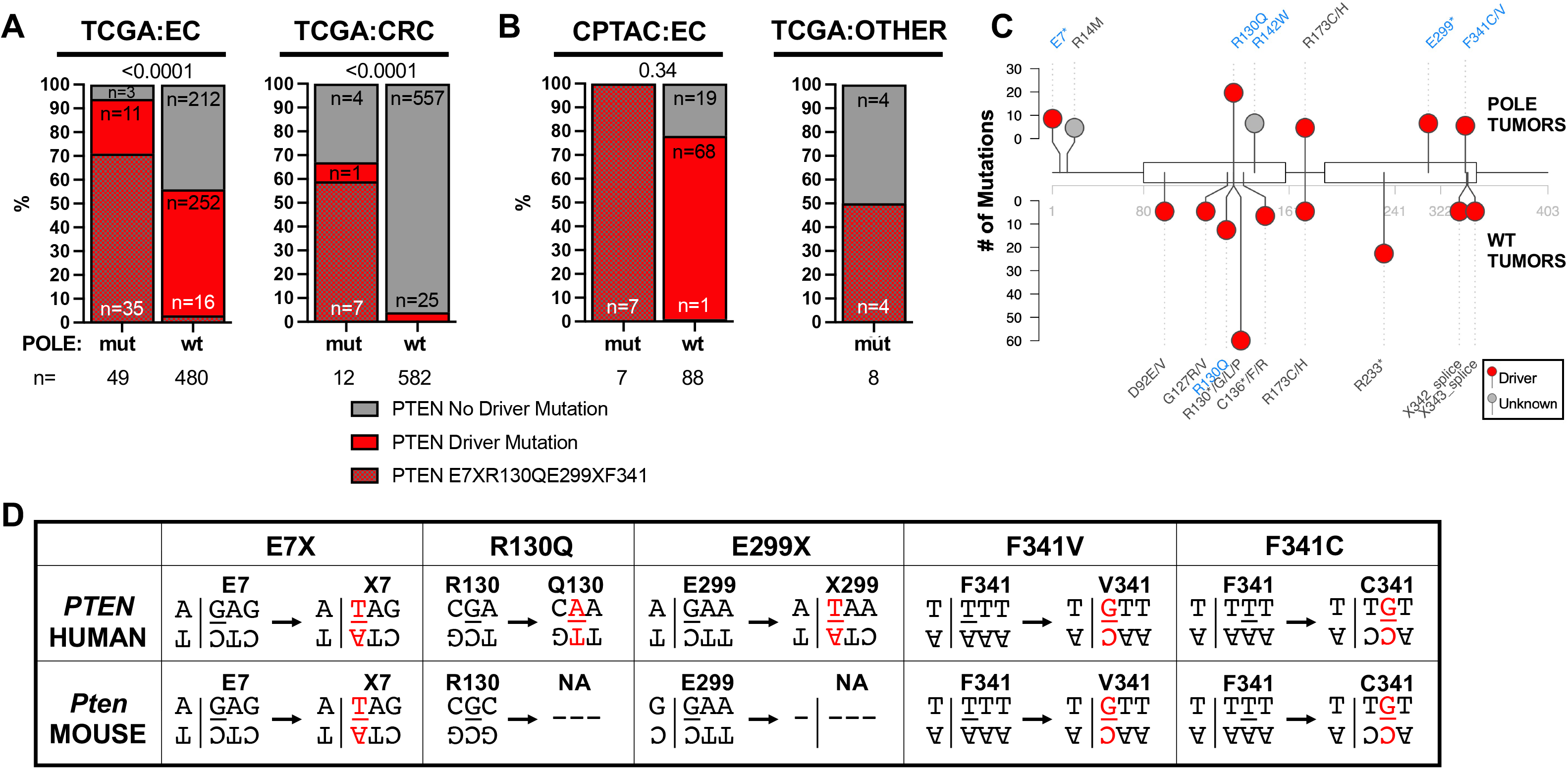
Driver mutations in *PTEN* are enriched in human and mouse POLE Tumors. **(A)** Frequencies of *PTEN* driver mutations are significantly enriched in human POLE mutant endometrial and colorectal tumors versus tumors with wild type POLE from the TCGA dataset. Fisher’s exact test was performed comparing all *PTEN* driver mutations, p-values indicated above stacked bars. **(B)** The frequencies of *PTEN* driver mutations are also elevated in POLE mutant tumors in the CPTAC endometrial dataset as well as in tumor types in TCGA that are not otherwise enriched in POLE mutations (All Other Tumors). Fisher’s exact test was performed comparing all *PTEN* driver mutations, p-values indicated above stacked bars. **(C)** Lollipop plot of *PTEN* mutations. Recurrent PTEN mutations (n>2) from POLE mutant (above) and POLE wild type tumors are plotted. Mutations occurring in POLE signature motifs are shown indicated in blue. **(D)** For each of the recurrent human PTEN driver mutations observed in POLE tumors, the flanking nucleotide context is shown (above) with the mutated base pair shown in red. The same flanking nucleotides are shown for the conserved residues in mouse Pten are also shown (below).

Of all *PTEN* driver mutations in POLE mutant tumors, we found significant enrichment in those occurring in POLE signature contexts, including recurrent E7X, R130Q, E299X, F341C and F341V *PTEN* mutations (Fig. 5B, C, Supp. Fig. 4, Supp Fig 5). R130 is critical for Pten catalytic activity ^36,37^ and F341V mutations have previously been shown to drive tumor formation in a mouse model ^38^. Together, these mutations accounted for 71% and 58% of all PTEN driver mutations in UCEC and CRC POLE mutant tumors, respectively, and also suggest that each was acquired due to aberrant DNA synthesis by the mutant DNA polymerase.

Based on the enrichment of *PTEN* mutations in human *POLE*-mutated tumors, we re-examined our *Pole*^+/*P286R*^ mouse tumors (this study and ^3^). We found that *Pten* mutations were enriched in *Pole*^+/*P286R*^ tumors from germline wild-type *Trp53* mice (50%, Supp. Table 3). These included multiple POLE signature mutations and several known driver mutations. We observed two occurrences of F341V *Pten* mutations in our mouse *Pole* mutant tumors. That this known PTEN driver mutation is recurrent in both human and mouse POLE mutant tumors strongly suggests a functional link between these two mutant alleles.

Analysis of protein levels in tumors can provide valuable information regarding the effects of gene mutations on protein function. We analyzed Pten protein levels in POLE mutant tumors reported in the National Cancer Institute’s Clinical Proteomic Tumor Analysis Consortium (CPTAC) endometrial proteogenomics study ^34^. Pten levels were lower in all tumors relative to normal tissues (Fig. 6A), consistent with *PTEN* mutation enrichment in endometrial tumors leading to reduced function. While *PTEN* nonsense mutations were weakly associated with lower Pten levels across all tumors, POLE mutant tumors with nonsense mutations had among the lowest Pten protein levels measured (Fig. 6B). And in the few samples that had matched normal and tumor tissues, POLE mutant tumors with nonsense mutations in *PTEN* were associated with the strongest decrease in Pten protein abundance (Fig. 6C). Interestingly, the PTEN R130Q mutation by itself saw no change in Pten protein levels, while the F341V mutations showed reduced, though not statistically significant, reductions in Pten protein levels (Fig. 6B). As previously reported for all tumors in the CPTAC set, p53 levels in POLE mutated tumors were not significantly different than adjacent normal tissue, even when accounting for p53 mutation status ^34^, tumor subtypes (Supp Fig. 6A) or within an individual with matched adjacent normal tissue (Supp. Fig. 6B).

**Figure 6.**
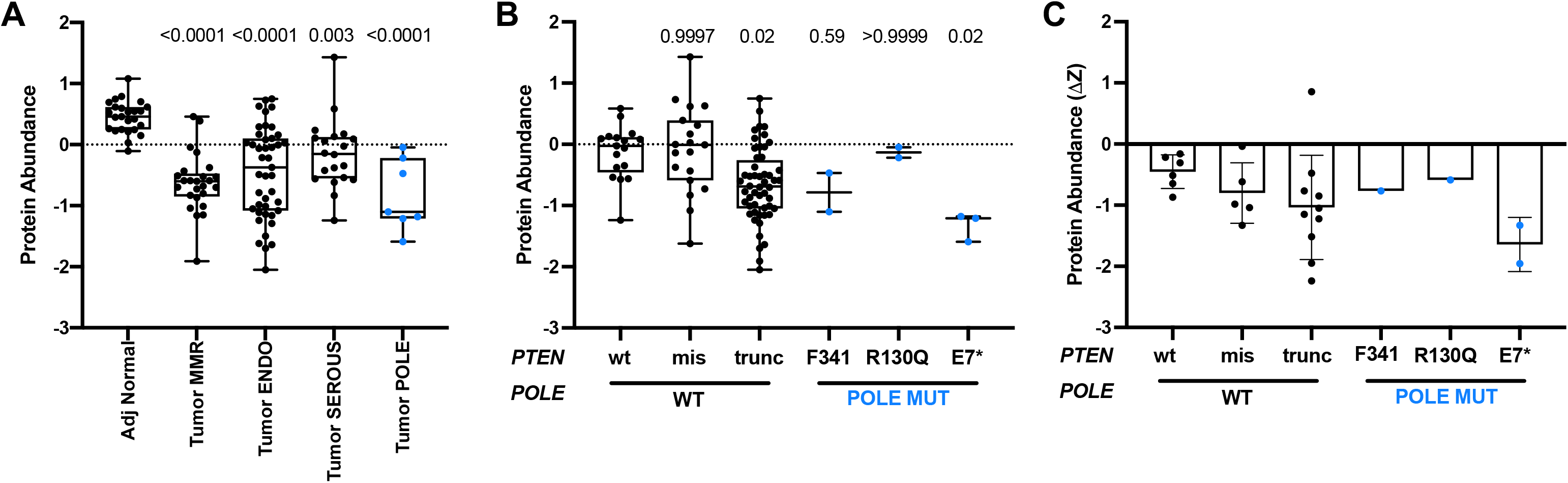
POLE-mediated *PTEN* mutations predict different effects on Pten protein abundance in human tumors. **(A)** *PTEN* protein abundance (z-score) of adjacent normal tissue and endometrial tumors, categorized into the four genomic subtypes, POLE, MSI, CNV-low (Endometroid-like (ENDO)), or CNV-high (Serous-like (SEROUS)) ^22, 39^. p-values indicated and determined by ANOVA. **(B)** Pten protein abundance (z-score) in tumors categorized by *PTEN* mutation status. Indicated p-values were determined by ANOVA. **(C)** Changes in individual tumor Pten protein abundance relative to matched normal (DZ) were plotted based on *PTEN* mutation status. p-values indicated and determined by ANOVA.

Strikingly, we found that the same *PTEN*-F341V recurrent mutation in human tumors also arose spontaneously in two different *Pole* mice. Engineered *Pten*^F341V/−^ mice develop tumors with a different tissue spectra than *Pten*^−/−^ mice ^38^. Caserta and colleagues made this mouse model based on observations that *PTEN*-F341V mutations were recurrent in human endometrial tumors. Re-analysis of the 10,967 tumors in the TCGA PanCancer studies shows that all 9 *PTEN*-F341V/C mutations occur in human tumors with *POLE* driver mutations. Leone *et al*. further described the *Pten-*F341V mutation in mice as contributing to tumor development through a non-canonical nuclear genome stability role for Pten. They showed evidence of reduced nuclear Pten protein coinciding with increased DNA damage in cells and tumors from these mice. We found γ-H2AX foci significantly enriched in a *Pole*^+/P286R^ tumor that spontaneously acquired a *Pten*-F341V mutation (Fig. 7), consistent with what was seen in a tumor from germline *Pten*^F341V/−^ mice, and consistent with a role for Pten in maintaining genome stability. Another mutation observed in our mouse cohort, *Pten*-D24L, is associated with lack of cytoplasm localization ^39^. This is supported by lack of staining in the tumor that harbors this mutation. The repeated co-occurrence of the *PTEN*-F341V mutation with mutations in *POLE* from tumors in both humans and mice suggests that both mutations may potentially cooperate in generating and subsequently failing to protect from genome destabilizing DNA damaging events or stress that contributes to tumorigenesis.

**Figure 7.**
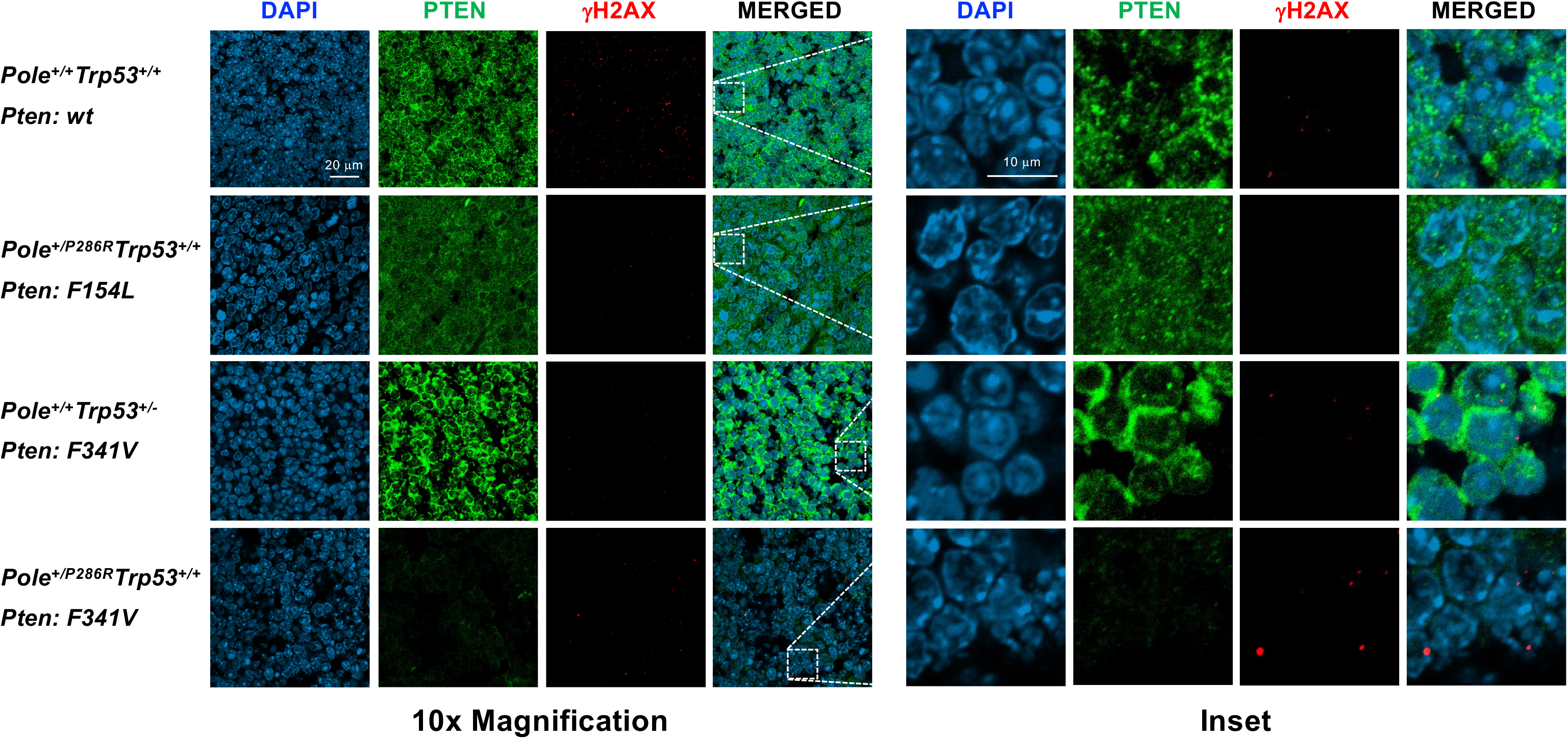
*Pten* mutations have differing effects on Pten subcellular localization and DNA damage levels in mouse tumors. Tumors from mice with the indicated Pole and Trp53 genotypes and somatic Pten mutations were processed and stained for DAPI (blue) and a-Pten (green) and a-gH2AX (red). Shown are representative confocal images at 10x magnification and zoomed in insets.

## DISCUSSION

Loss of p53 function, which rescues the viability of many mutant alleles, including *Pole4*^−/−^, failed to render *Pole*^P286R/P286R^ mice viable. Inactivation of one *Trp53* allele accelerated tumorigenesis via POLE-mediated mutation in the remaining *Trp53* allele. Data collected from cells suggests that the mutant Pole-P286R polymerase activates p53, suggesting possible selective pressure towards inactivating p53 via mutation. Mutagenesis *per se* is unlikely to be the primary cause of p53 activation, as we observed no difference in mutation burden or signature between mouse tumors with or without p53 mutations. Instead, POLE mutant tumors are significantly biased towards mutations in *PTEN*. The recurrence in both human and mouse tumors of a *PTEN* mutation that specifically reduces nuclear Pten abundance and fails to guard against genome instability suggests involvement of mutant Pole in the replication stress that is normally surveilled by Pten.

As the most frequently mutated gene in cancer, loss of p53 function, whether through mutational inactivation or inhibition, has long been known to accelerate tumor development ^40,41^. Despite this vast knowledge, it was unclear until recently to what extent loss of p53 function impacted base pair substitutions and indels. A comprehensive analysis of 10,000 whole exomes from tumors in TCGA found a minimal effect (2.2-fold) of *TP53* mutations on overall tumor mutation burden and no change in mutational signatures ^42^. The lack of unique mutational signatures in p53 mutant tumors led to the idea that loss of p53 function is unlikely to cause *de novo* mutagenesis, but instead permits the accumulation of mutations via other pathways ^23,42^. We find that this remains true in POLE mutant tumors where there is no difference in mutation burden or signature between those tumors with or without p53 driver mutations.

Coupled with this absence of an effect on mutagenesis, the relative lack of *TP53* and *Trp53* mutations in human and mouse tumors led us to explore other possible cancer driver gene mutations that might play roles in driving POLE mutant tumor development. This type of analysis is complicated by the fact that since POLE mutant tumors acquire so many mutations, essentially every gene has a mutation. Thus, discerning driver mutations from passengers presents certain challenges. Our previously engineered *Pole*^+/P286R^ mice with wild type *Trp53* provided an excellent opportunity to address this complication.

Potential cancer driver genes contributing to POLE mutant tumorigenesis would be predicted to share several characteristics. Mutations should be enriched in the same gene in both human and mouse POLE mutant tumors. Mutations should be enriched either in the same residue, or, if homology is low, then should be enriched in functionally conserved residues or motifs. Mutations should occur repeatedly, but not necessarily exclusively, in POLE signature motifs. Finally, ideally, there should be evidence indicating that those mutations disrupt normal gene product function. Mutations satisfying all these criteria in both mouse and human POLE mutant tumors would provide corroborating evidence supporting the idea that those genes are in fact cancer driver genes.

Mutations in *PTEN* satisfy all these criteria and further hint at possible mechanisms of how Pten dysfunction may interact with the mutant polymerase to promote tumor development. First, *PTEN* mutations are significantly enriched in both mouse and human POLE mutant tumors. More than 90% of TCGA endometrial tumors with mutations in *POLE* also have a known driver mutation in *PTEN* (Fig. 5A), and enrichment is also seen in colorectal cancers. POLE mutant tumors with *PTEN* mutations also occur sporadically in other tumor types.

Several specific *PTEN* mutations occur in both human and mouse POLE mutant tumors, further supporting the idea that the PTEN and POLE pathways are interacting. Once such mutation, F341V/C, occurred twice in our mouse tumors and is recurrent in human POLE mutant tumors, likely due to a combination of its occurring in a POLE signature context and its unique effects on Pten function discussed below. The PTEN-F154, Y68 and D24 mutations found in our POLE mouse tumors are also recurrent in human tumors with wild type POLE, suggesting that these are not simply passenger mutations (Supp. Table 4).

Mutational signatures can also explain both the occurrence of a hotspot mutation as well as differences between species abundance. The strongest *PTEN* hotspot mutation (R130Q) and highly recurrent nonsense mutation (E299X) observed in human POLE mutant tumors both have different sequence contexts in mice that preclude their mutation in mouse tumors (Fig. 5C), thus explaining their absence in our mouse tumors. The E7X mutation, however, is recurrent in human tumors yet absent in mice. The simplest explanation is that our tumor sample size is too small.

The functional effects of mutations in PTEN have been examined extensively using many different assays. With the exception of F154L, all *Pten* mutations we observed in mouse POLE mutant tumors are either known to cause loss of function ^38,43^, or are immediately adjacent to a known susceptible residue (I122L). Just looking at known *PTEN* driver mutations, we find that nonsense mutations significantly reduce the amount of Pten present in actual human POLE mutant tumors (Figs. 5 and 6). The unchanged levels of Pten-R130Q and only modestly changed Pten-F341V/C suggest that these missense mutations may have novel functions critical to enhancing POLE mutant tumor development.

How might Pten defects help promote POLE mutant tumor development? The simplest explanation is through canonical loss of Pten and subsequent PI3K-AKT activation of downstream targets that enhance cellular growth. This is the likely situation in tumors with PTEN nonsense mutations. However, several lines of evidence suggest that the primary mechanism might be more complicated, possibly through increased replication stress tolerance. Pten-F341 is a recurrent mutation site in both mouse and human POLE mutant tumors, yet *Pten*^F341V/F341V^ mice develop tumors while retaining normal Akt activation and embryonic development ^38^. Heterozygous mutations at F341 are sufficient to drive tumorigenesis as *Pten*^+/F341V^ mice develop tumors in a number of different organs, including the thymus. Even more intriguing is the observation that Pten is depleted specifically from nuclei from *Pten*^F341V^ mice, which also coincides with increased incidence of γ-H2AX foci. These results are consistent with other work showing that Pten has nuclear roles in suppressing genome instability, chromatin compaction, chromosome condensation, checkpoint control and replisome stability ^44,45^. Pten has both phosphatase-dependent and -independent activities that are critical in promoting both normal replisome assembly as well as stalled fork protection and reassembly, particularly through targets including Mcm2, Rpa1 and Rad51 ^46–48^.

Nonsense mutations (e.g. E7X, E299X) and other missense mutations that destabilize the Pten protein (e.g. R173C/H, F341C/V) would certainly lose the ability to regulate the PI3K-AKT pathway. But they would also have the same loss of function effects on nuclear Pten activities. The remaining *PTEN* hotspot mutation in POLE mutant tumors is R130Q. The R130G mutation makes a stable protein in cells and both R130G and R130Q show increased p-Akt in glioblastomas and endometrial tumors, suggesting loss of lipid phosphatase activity in POLE mutant tumors ^49^. However, mutations at R130 have been proposed to act by suppressing the activity of the wild type partner in a mixed heterodimer in a dominant negative fashion. Since the effects of R130 dominant negative mutation have not yet been tested on nuclear Pten function, it is tempting to speculate that the enriched PTEN-R130Q is specifically helping facilitate POLE mutant tumor development via interfering with a nuclear genome stabilizing activity. Loss of PI3K-AKT regulation would serve as an added unrelated tumorigenic boost.

Our observations that POLE mutant tumor formation in *Trp53*^+/−^ mice is accelerated, likely via *Trp53* LOH, is also consistent with mutant *Pole* alleles triggering replisome dysfunction. We showed that cells expressing the mutant polymerase enzyme activate p53. Both p53 and Pten could be involved in suppressing possible resulting genome instability. Normally two hits would be required to inactivate p53 and relieve this suppression, while a single hit to Pten would begin to relieve this suppression. The observation that mutant Pol ε has hyperpolymerization characteristics ^50^ is particularly intriguing in light of our results in POLE mutant tumors.

Taken together, we propose that development of tumors with mutations in *POLE* is dependent on two features. The elevated mutagenesis is required to supply the loss of function mutations in specific genes, but is by itself is insufficient to drive tumor formation. Also required is a loss of genome instability suppression activity, possibly involving replisomes that have become dysfunctional via engaging the mutant Pol ɛ.

## MATERIALS AND METHODS

### Pole-P286R and Trp53 Genotyping Strategy

Tail gDNA was extracted and PCR amplified. PCR products were directly treated with DdeI restriction enzyme (NEB) per product protocol and run on agarose gel as previously described ^3^. The *Pole* locus was PCR amplified using the following primers:

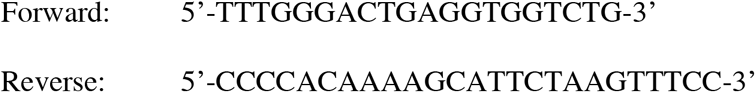

The *Trp53* locus was PCR amplified using the following primers:

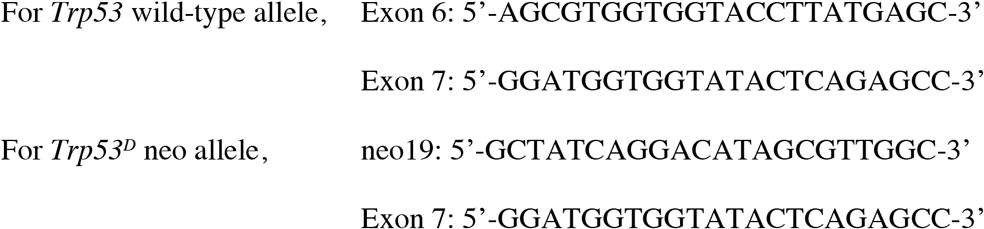

### Survival Analysis

Mice were housed and maintained at the Tulane University Health Science Center vivarium and were under care and supervision of staff veterinarian and trained animal care and veterinary technicians. All animals were monitored closely and sacrificed by CO_2_ inhalation if the following criteria were observed: ulcerating tumor, visible tumor >1 cm, moribund, visible weight loss (>10%), etc.

### Targeted Panel and Whole Genome Sequencing of Tumors

Genomic DNA from tumor and tail of mice were extracted by phenol/chloroform extraction. Samples were sent to the Beijing Genomics Institute Americas for sequencing. Agilent SureSelectXT Mouse All Exon Kit was used for enrichment. Illumina HiSeq4000 was used for paired-end sequencing.

### Variant Detection From Next Generation Sequencing Data

FASTQ files were aligned to the reference genome using BWA-MEM 0.7.12, and PicardTools 1.133 to sort the BAM file and mark duplicates. Variant calling was completed as described ^3^.

### Mouse Embryonic Fibroblast Generation and Population Doubling Level Experiments

Mouse embryonic fibroblasts (MEFs) were derived from timed mating of double heterozygous crosses (*Pole*^+/*P286R*^;*Trp53*^+/−^ × *Pole*^+/*P286R*^;*Trp53*^+/−^) at E12.5-13.5 from viable *Pole* wild-type, heterozygous and homozygous embryos. Genomic DNA from embryo heads were used for genotyping, while the remaining embryo was minced, trypsinized and plated for cell culture under aseptic conditions in 10 cm plates. After 24-48 hours, cells were counted.

For population doubling level (PDL) measurement, 1.9 × 10^5^ cells were seeded in 6-well plates in triplicate and grown at 37°C in 5% CO_2_. After 3 days, each triplicate was harvested individually and live cells were counted using a Countess Automated Cell Counter (Invitrogen). This procedure was repeated every 3 days until the cells were immortalized as per NIH3T3 protocol. PDL was calculated as follows:

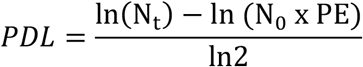

where Nt = number of viable cells counted after passage; N0 = number of cells seeded prior to passage; PE = plating efficiency. Passage 0 was the initial seeding from the 10 cm plate into 6-well plates.

### Expression of p53 Targets

Total RNA was extracted using Invitrogen TRIzol Reagent (Cat. #15596026) and chloroform as per product protocol. RNA was precipitated in isopropanol and resuspended in DEPC water. RNA was immediately made into cDNA (First-Strand cDNA Synthesis Kit, GE). Real time PCR was performed using QuantStudio 6 (ThermoFisher Scientific). Expression of targets was normalized to TBP. The following primers were used:

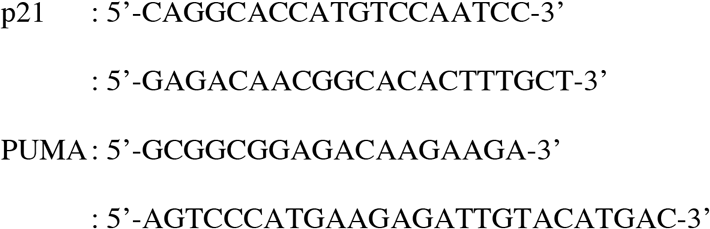

### Immunofluorescent Staining

Mouse thymus tissues were harvested and fixed in 10% formalin by standard methods. Parafin-embedded tissues were sectioned, mounted in xylene, boiled in citrate buffer, then stained with rabbit anti-Pten (Cell Signaling Technologies Cat.# 9559T), mouse anti-phospho Histone-H2A.X (Ser139) (EMP Millipore Cat.# 05-636) and DAPI (Invitrogen Cat.# D1306).

### Driver Gene Mutation Enrichment Analysis in Human Cancer Cohorts

Frequency of TP53 and PTEN driver mutations in POLE mutant and POLE wild type tumors were counted, and proportions were analyzed via Fisher’s exact test to account for small sample sizes.

### CPTAC Endometrial Carcinoma ^34^ Protein Analysis

Individual sample protein abundance plotted based on reported z-score per Dou, et al. Individual ΔZ calculated: ΔZ = Z_Tumor_ – Z_Adjacent Normal Tissue_ if adjacent normal tissue protein analysis was reported.

