## Supplementary Figures for "Mouse model and human patient data suggest critical roles for Pten and p53 in suppressing POLE mutant tumor development"

### Supp Fig 1

**A**

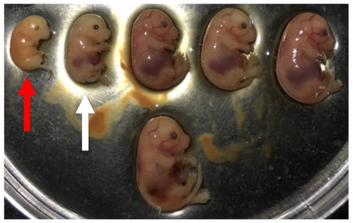

**B**

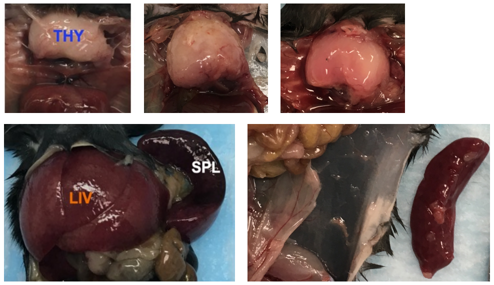


**Supplementary Figure 1. A *Pole*$^{P286R/P286R}$ embryo is severely underdeveloped compared to littermates. (A)** Image of litter of double heterozygous cross. Exact embryonic day is unknown. Red arrow indicates *Pole*$^{P286R/P286R}$ fetus. Genotypes from left to right: *Pole*$^{P286R/P286R}$*;Trp53*$^{+/-}$ *(red arrow), Pole*$^{+/P286R}$*;Trp53*$^{-/-}$ *(white arrow), Pole*$^{+/P286R}$*;Trp53*$^{+/-}$*, Pole*$^{+/P286R}$*;Trp53*$^{+/-}$*, Pole*$^{+/P286R}$*;Trp53*$^{+/-}$*, Pole*$^{+/+}$*;Trp53*$^{+/-}$. **(B)** *Pole*-mutated mice primarily develop lymphomas. Images are representative of lymphomas observed in Pole$^{+/P286R}$Trp53$^{+/-}$ mice. Frequency noted in Supp Table 2.

### Supp Fig 2

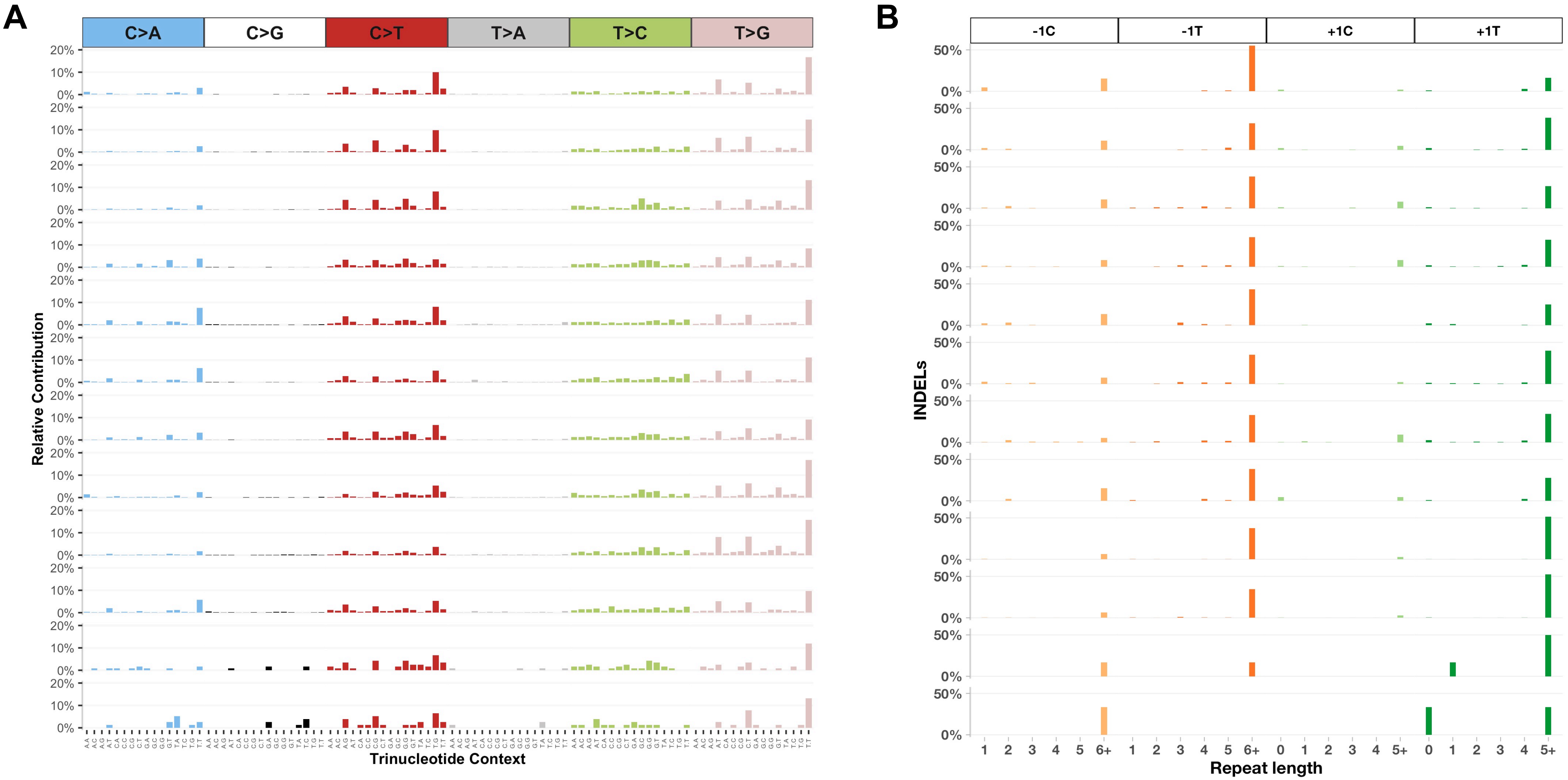


**Supplementary Figure 2. *Pole*$^{+/P286R}$*;Trp53*$^{+/-}$ tumors show evidence of POLE SBS 10a, 10b, 28 as well as +1 insertions in homopolymeric T runs. (A)** Mutation spectrum of *Pole*$^{+/P286R}$*;Trp53*$^{+/-}$ thymic tumors (top 2 rows) and *Pole*$^{+/P286R}$*;Trp53*$^{+/-}$ splenic tumors (bottom two rows). **(B)** Insertion/deletion spectrum of *Pole*$^{+/P286R}$*;Trp53*$^{+/-}$ thymic tumors (top 2 rows) and *Pole*$^{+/P286R}$*;Trp53*$^{+/-}$ splenic tumors (bottom two rows).

#### Supp Fig 3

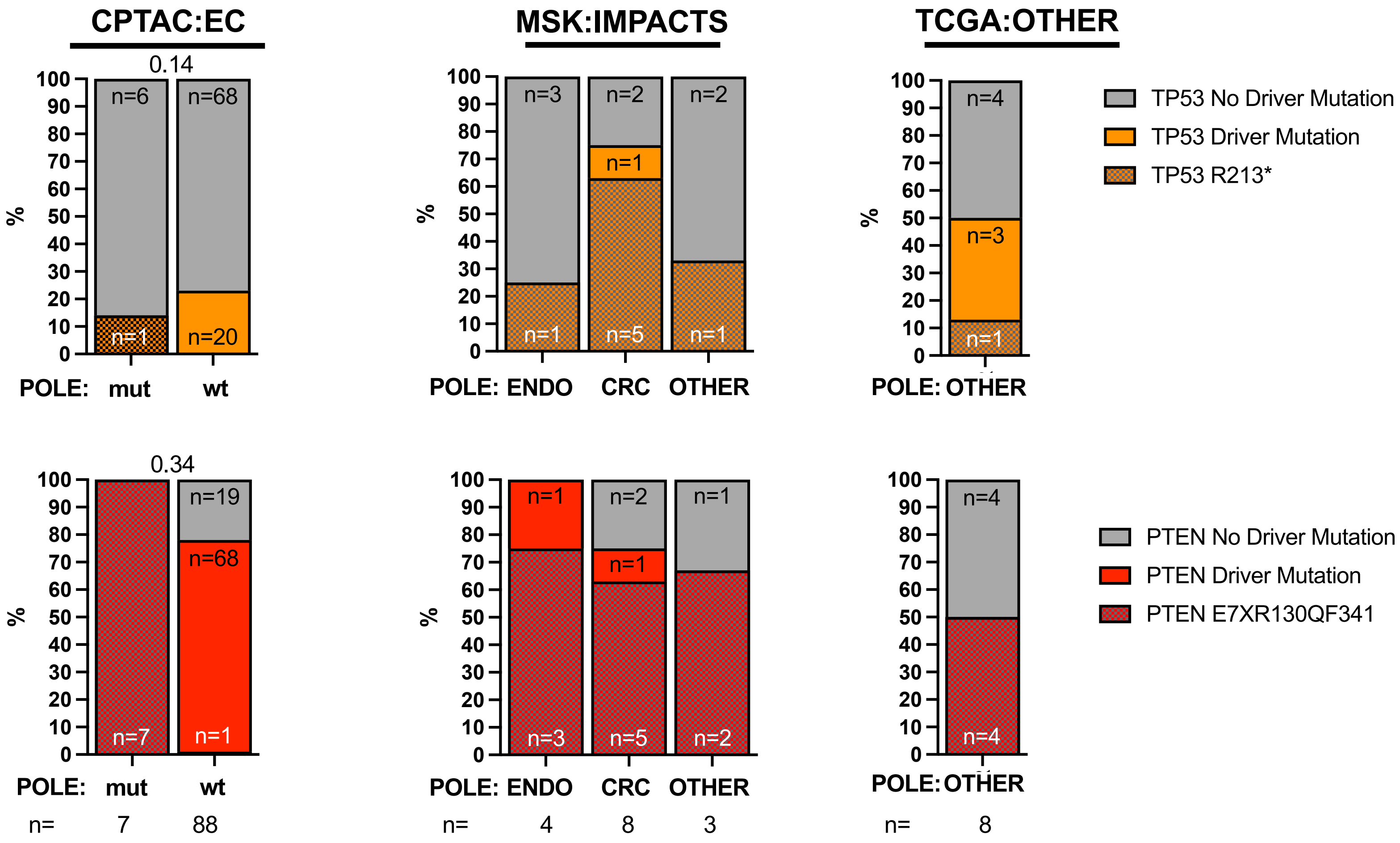


**Supp Fig 3. *TP53* (orange) and *PTEN* (red) driver mutations in various cohorts from POLE mutant or POLE wt tumors.** Shown are the per cent of total and number of each in the following cohorts: Endometrial CPTAC; Endometrial and Colorectal tumors from MSK:Impacts cohort; POLE mutant tumors from TCGA non-EC/CRC cohort.

### Supp Fig 4

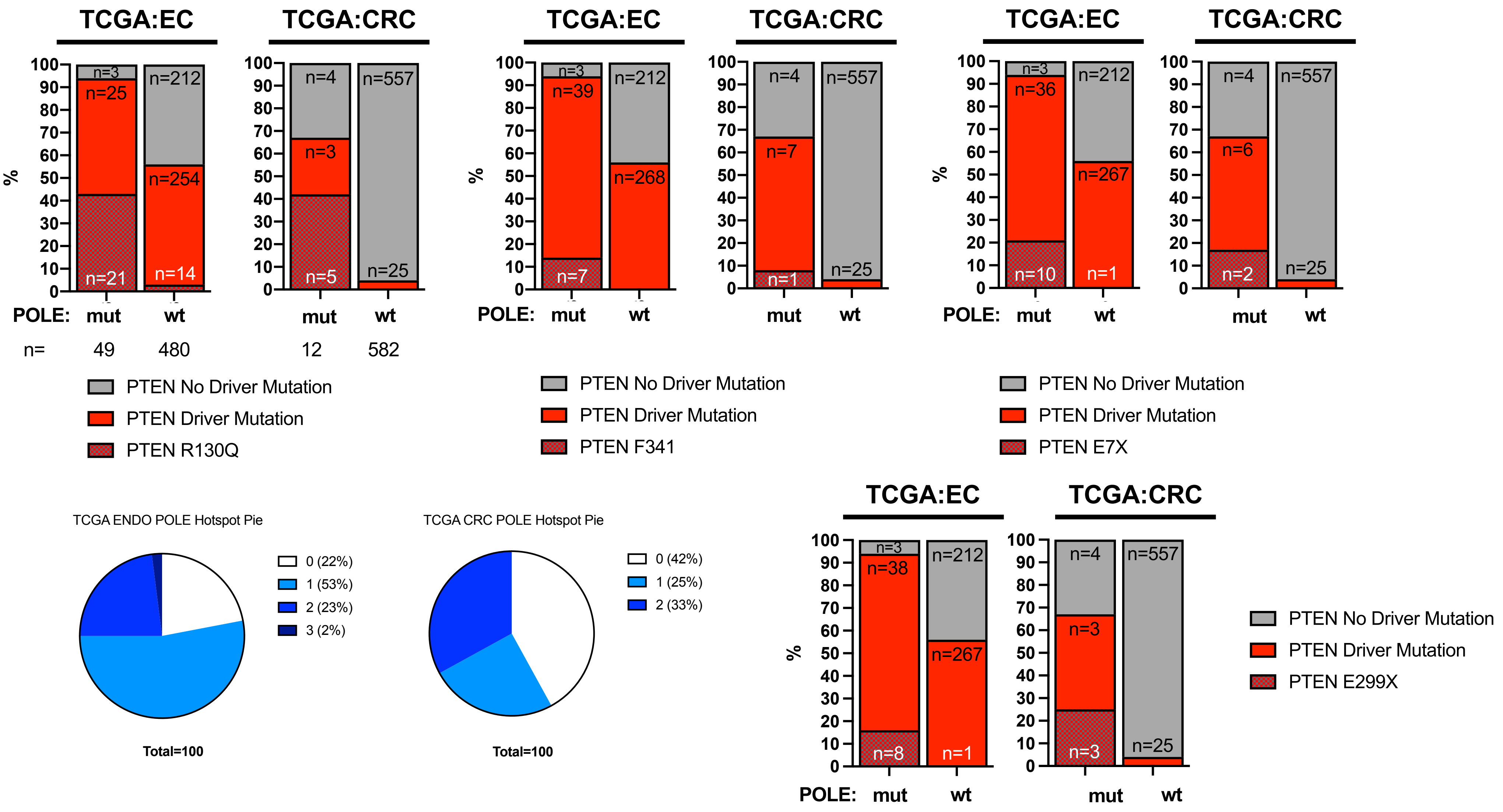
TCGA:EC
TCGA:CRC
%
100
90
80
70
60
50
40
30
20
10
0
n=3
n=25
n=21
n=212
n=254
n=14
n=4
n=3
n=5
n=557
n=25
POLE: mut wt
mut wt
n= 49 480 12 582
PTEN No Driver Mutation
PTEN Driver Mutation
PTEN R130Q
TCGA:EC
TCGA:CRC
n=3
n=39
n=7
n=212
n=268
n=4
n=7
n=1
n=557
n=25
POLE: mut wt
mut wt
PTEN No Driver Mutation
PTEN Driver Mutation
PTEN F341
TCGA:EC
TCGA:CRC
n=3
n=36
n=10
n=212
n=267
n=1
n=4
n=6
n=2
n=557
n=25
POLE: mut wt
mut wt
PTEN No Driver Mutation
PTEN Driver Mutation
PTEN E7X
TCGA ENDO POLE Hotspot Pie
0 (22%)
1 (53%)
2 (23%)
3 (2%)
Total=100
TCGA CRC POLE Hotspot Pie
0 (42%)
1 (25%)
2 (33%)
Total=100
TCGA:EC
TCGA:CRC
n=3
n=38
n=8
n=212
n=267
n=1
n=4
n=3
n=3
n=557
n=25
POLE: mut wt
mut wt
PTEN No Driver Mutation
PTEN Driver Mutation
PTEN E299X

#### Supp Fig 5

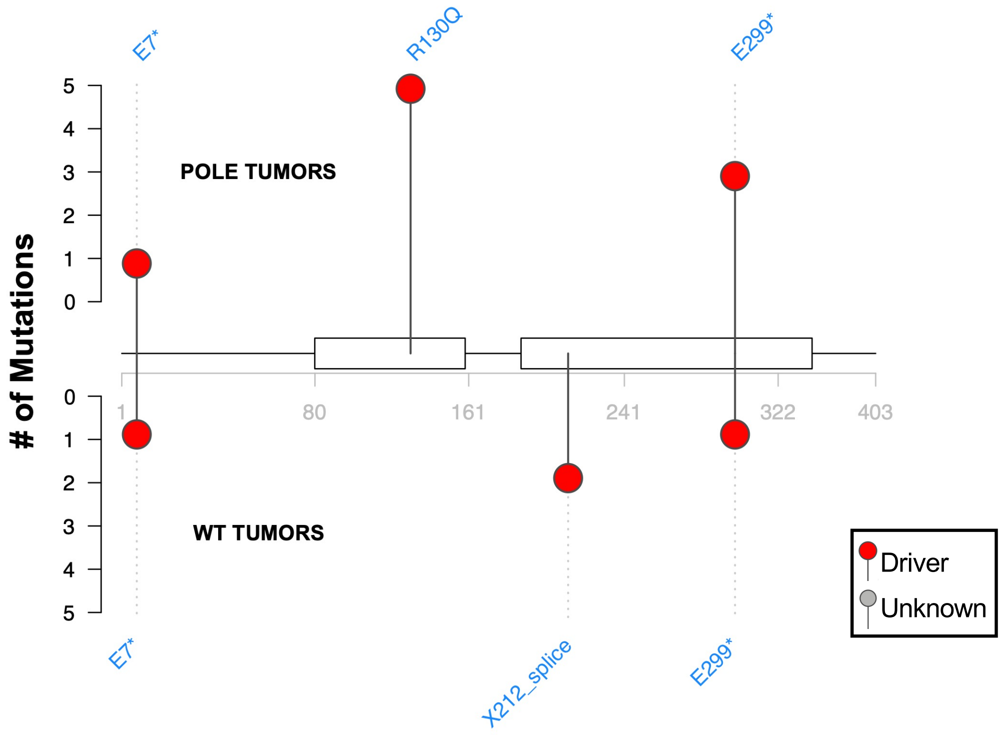


**Supplemental Figure 5. Driver mutations in *PTEN* in TCGA CRC POLE mutant and POLE WT Tumors.** Lollipop plot showing the number of driver (red) and unknown significance (gray) *PTEN* mutations.

### Supp Fig 6

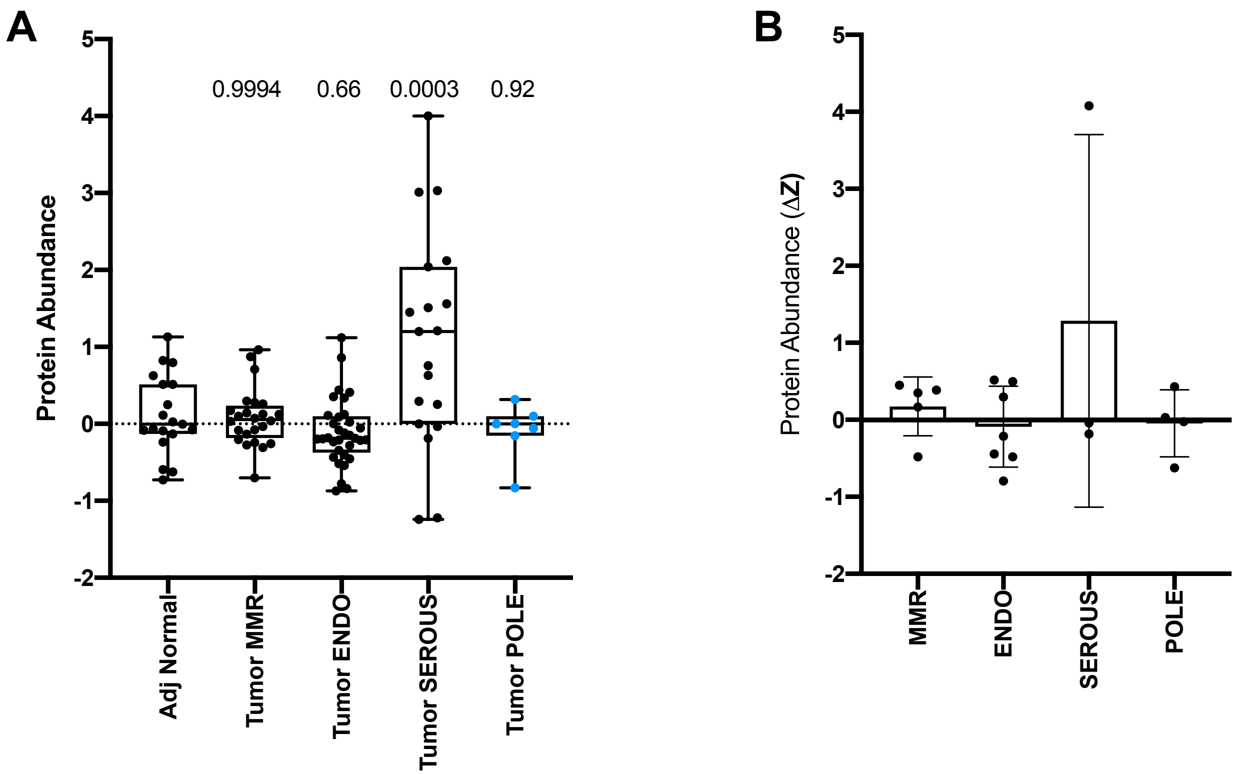


**Supplementary Figure 6.** ***TP53* protein abundance in CPTAC endometrial cancer cohort in tumor subtypes. (A)** *TP53* protein abundance (z-score) of adjacent normal tissue and endometrial tumors, categorized into the four genomic subtypes, POLE, MSI, CNV-low (Endometroid-like (ENDO)), or CNV-high (Serous-like (SEROUS)) [22, 39]. p-values indicated and determined by ANOVA. **(B)** Z-score of tumor minus z-score of adjacent normal tissue (delta Z (ΔZ)) of individual tumor categorized categorized into the four genomic subtypes, POLE, MSI, ENDO, SEROUS.
